# Reprogramming of three-dimensional microenvironments for *in vitro* hair follicle induction

**DOI:** 10.1101/2022.06.13.495917

**Authors:** Tatsuto Kageyama, Akihiro Shimizu, Riki Anakama, Rikuma Nakajima, Kohei Suzuki, Yusuke Okubo, Junji Fukuda

**Affiliations:** Faculty of Engineering, Yokohama National University, 79-5 Tokiwadai, Hodogaya-ku, Yokohama, Kanagawa 240-8501, Japan; Kanagawa Institute of Industrial Science and Technology, 3-2-1 Sakado Takatsu-ku, Kawasaki, Kanagawa 213-0012, Japan; Japan Science and Technology Agency (JST)-PRESTO, 4-1-8 Honcho, Kawaguchi, Saitama, 332-0012, Japan; Division of Cellular & Molecular Toxicology, Center for Biological Safety & Research, National Institute of Health Sciences, 3-25-26 Tono-machi, Kawasaki-ku, Kawasaki, Kanagawa 210-9501, Japan

**Keywords:** Hair follicle regeneration, Hair follicle organoid, Self-organization, Epithelial-mesenchymal interaction

## Abstract

During embryonic development, reciprocal interactions between epidermal and mesenchymal layers trigger hair follicle morphogenesis. This study revealed that microenvironmental reprogramming via control over these interactions enabled hair follicle induction *in vitro*. A key approach is to modulate spatial distributions of epithelial and mesenchymal cells in their spontaneous organization. The *de novo* hair follicles with typical morphological features emerged in aggregates of the two cell types, termed hair follicloids, and hair shafts sprouted with near 100% efficiency *in vitro*. The hair shaft length reached ∼3 mm in culture. Typical trichogenic signaling pathways were upregulated in hair follicloids. Owing to replication of hair follicle morphogenesis *in vitro*, production and transportation of melanosomes were also monitored in the hair bulb region. This *in vitro* hair follicle model might be valuable for better understanding hair follicle induction, for evaluating hair growth as well as the inhibition of hair growth by drugs, and modeling gray hairs in a well-defined environment.

**Teaser:** In tissue morphogenesis, different types of cells harmonize in a pre-programmed manner using messenger systems such as epithelial-mesenchymal interactions. Organoids are a promising tool to elucidate such mechanisms on a molecular level. This work describes a strategy for reprograming three-dimensional microenvironments to trigger the initiation of *in vitro* regeneration of hair follicle organoids. Hair follicle organoids generated fully matured hair follicles, enabling the monitoring of hair follicle morphogenesis *in vitro* and determination of signaling pathways involved in early hair follicle morphogenesis. The principles uncovered herein may be relevant to other organ systems and will contribute to our understanding of developmental phenomena in physiological and pathological processes, eventually opening up new research avenues for the development of new treatment strategies.

## Introduction

During the early developmental stages, epithelial-mesenchymal interactions (EMIs) trigger the development of various tissues and organs through specific signaling pathways and transcription factors. For example, EMIs drive stomach specification (*1*), duct patterning of the kidney (*2*) and lung (*3*), and bud formation of the lacrimal gland (*4*) and teeth (*5*). Additionally, the development of the hair follicle is coordinated by EMIs. However, the hair follicle is a unique mini-organ that repetitively and dynamically undergoes cyclic regeneration postnatally (*6, 7*). Thus, it has been studied as a representative model to understand underlying mechanisms responsible for EMIs in complex development (*8*).

Over the last several decades, some crucial mechanisms of EMIs, including the effect of *Wnt* signaling on hair follicle morphogenesis (*9*), have been elucidated using animal models. Although knockout and knockdown mouse models can be used to identify key genes and signals related to hair follicle development based on the appearance of body hairs, fully elucidating molecular mechanisms for EMIs remains challenging due to the crowded *in vivo* environment. *In vitro* three-dimensional and organoid culture approaches for various tissues and organs have recently received widespread attention because these *in vitro* approaches simplify the model system with close microscopic observation. Hair follicle germ (HFG)-like aggregates were reconstructed in culture using dissociated embryonic epithelial and mesenchymal cells (*10*). When transplanted into the skin of nude mice, HFGs generated *de novo* hair follicles, implicating that HFGs possess hair neogenesis capability. However, it remains challenging to induce hair neogenesis in *in vitro* culture, including the generation of matured hair follicles (e.g., dermal papilla and the bulge region) and the extension of long hair shafts (*11*). Skin organoids including hair follicles and sebaceous glands were generated *in vitro* using mouse pluripotent stem cells through self-organization of epidermal and dermal layers (*12*). More recently, human induced pluripotent stem cells were used to generate skin organoids with appendages including hair follicles in 4–5 months of culture (*13*). The skin organoids were shown to be equivalent to the embryonic skin in development and mimicked normal hair folliculogenesis. However, skin organoid culture requires multiple differentiation steps starting with ectoderm induction on the surface of cell aggregates, takes a relatively long period of time, contains unexpectedly differentiated cells and their secreted proteins, and is limited to relatively short lengths of hair shafts.

We have reported an approach for engineering HFG-like aggregates *in vitro* through the self-organization of embryonic dorsal epithelial and mesenchymal cells in a lab-made microwell array plate (*14*). In this approach, the two types of rodent cells initially formed a single aggregate where the two cell types were distributed randomly, and subsequently spatially separated and formed HFG-like structures during 3 days of culture. When transplanted into the skin of nude mice, the HFGs efficiently generated *de novo* hair follicles in a few weeks and represented normal hair cycles. Furthermore, HFGs generated hair follicles and shafts *in vitro* in a long-term culture, whereas the appearance of hair follicles was quite rare.

In this study, we demonstrate that hair follicles are generated from almost all the HFGs (∼100% of HFGs) by reprogramming cell microenvironments through the modulation of EMIs associated with the spatial distribution of epithelial and mesenchymal cells. Using the *in vitro* hair follicle neogenesis model, we examined the developmental mechanisms and the signaling pathways involved in early hair follicle morphogenesis. Furthermore, taking advantage of this *in vitro* system, we continuously monitored the production and transportation of melanosomes in hair bulbs. This model may provide a platform for better understanding hair follicle neogenesis and pigmentation.

## Results

### Control over the spatial distribution of cells for hair follicle neogenesis in vitro

The formation of HFGs during development triggers hair follicle induction through EMIs, which motivates researchers to engineer tissue grafts for hair regeneration (*10, 11*). We have previously reported that HFG-like aggregates prepared through self-organization can generate hair follicles when transplanted (*14*). In the present study, we found that a hair shaft-like protruding fiber appeared *in vitro* from the dumbbell-shape HFGs after 6 days of culture (Supplementary Fig. 1a, b). The fibers became ∼200 µm in length at 23 days of culture, which possessed typical morphological features such as the hair cortex and micro-fibrils (Supplementary Fig. 1c). Although the sprouting occurred only in < 1% of HFGs (1/300 HFGs), this matured hair follicle formation encouraged us to seek an approach for *in vitro* hair follicle neogenesis. To reproduce this phenomenon more efficiently, we investigated key factors that trigger the hair follicle sprouting, including types of basal media, growth factors, ratios of the two cell types, and total cell numbers. Among them, fibroblast growth factor-2 (FGF2) was effective for HFGs to increase the expression of two trichogenic genes, *versican* and *Tgfβ2* (Supplementary Fig. 1d), which is consistent with a previous study (*15*). FGF2 slightly improved the frequency of appearance of hair follicle sprouting (6/300 HFGs). In the present study, we defined the sprouting efficiency as the average number of hair follicle sprouting per HFG, which was improved from 0.003 to 0.02 hairs/HFG by the addition of FGF2 (Supplementary Fig. 1e).

A key finding of this study was that the supplement of Matrigel (or other extracellular matrices, ECMs, as describe later) significantly improved the sprouting efficiency to ∼1.0 (∼300/300 HFGs) (Fig. 1). Matrigel has widely been used for organoid culture of various tissue types by encapsulating stem cells in its hydrogel form (*16*). However, in the present study, Matrigel was supplemented at a sufficiently low concentration (2v/v%) and kept at 4 °C even after cell seeding for at least 30 min so as not to form a hydrogel and not to prevent cell aggregation, since cell-cell contact between epithelial and mesenchymal cells is critical to induce EMIs. Note that the supplement of a low concentration of Matrigel is not novel and has been reported in research on organoids, including inner ear and skin organoids (*12, 17*), and the cooled cell aggregation processes were vital for our system. In this culture, the two types of cells formed an aggregate and spontaneously separated from each other in the aggregate as expected, but their behaviors were completely different from those without supplement of Matrigel. The cells did not form a dumbbell-shape aggregate but formed an aggregate composed of a core of epithelial cells surrounded by mesenchymal cells (Fig. 2A). Interestingly, multiple pigmented portions emerged in the single aggregates at 2 days of culture, from which hair follicles sprouted (Fig. 2A, arrows, Supplementary Movie 1). This hair follicle sprouting model was termed as hair follicloid hereafter. The hair follicloids at 2 days of culture were embedded in 100v/v% Matrigel in gel form. Without the hydrogel encapsulation, the sprouted hair follicles remained around periphery of the hair follicloids (data not shown). However, with this two-step Matrigel supplement culture, the hair shafts extended straightly due to the hydrogel supports, which is consistent with a previous report on skin organoids (*12*). The length of hair shafts reached ∼3 mm at 23 days (Fig. 2B).

**Fig. 1.**
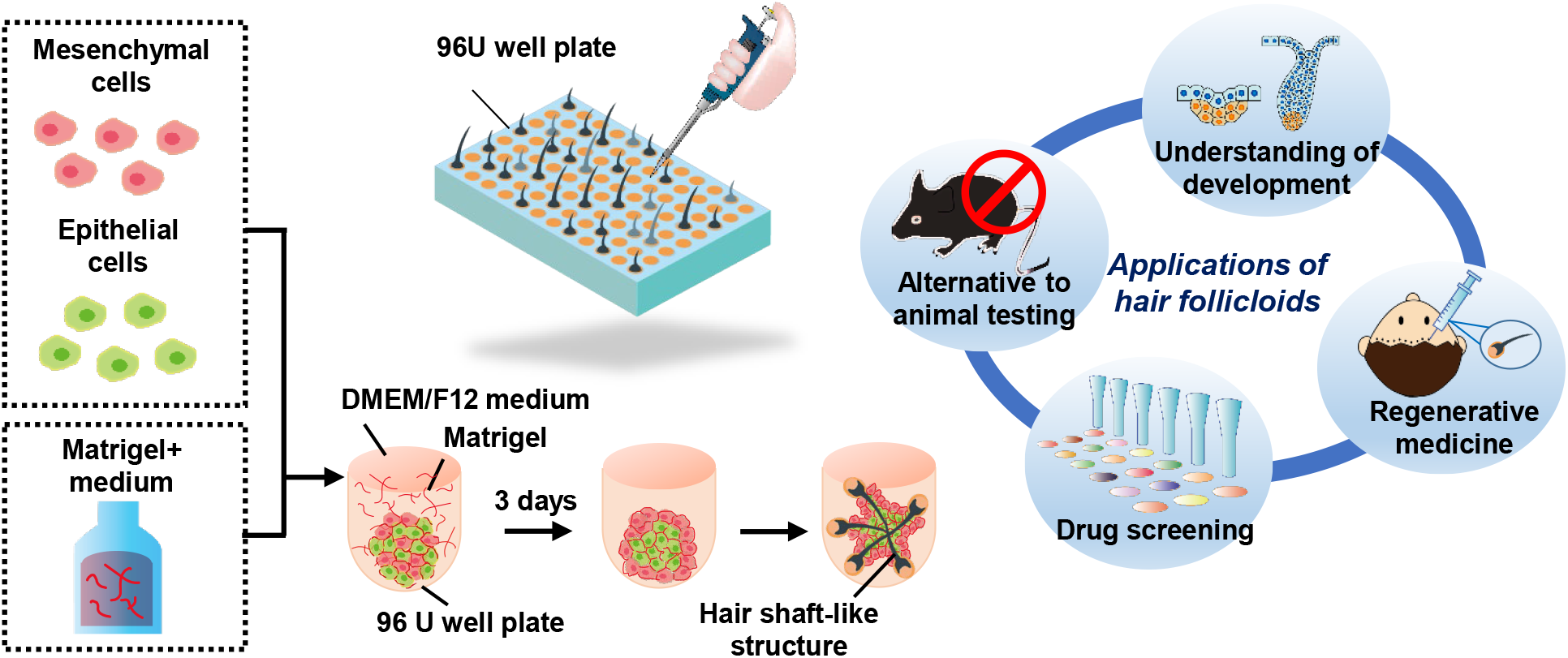
Schematic representation of the preparation of hair follicloids. Hair follicloids formed through the self-organization of embryonic epithelial and mesenchymal cells generated hair shafts *in vitro* with a high level of efficiency, and are potentially beneficial for several applications including understanding of development and pathology, alternative to animal testing, drug screening, and regeneration medicine.

**Fig. 2.**
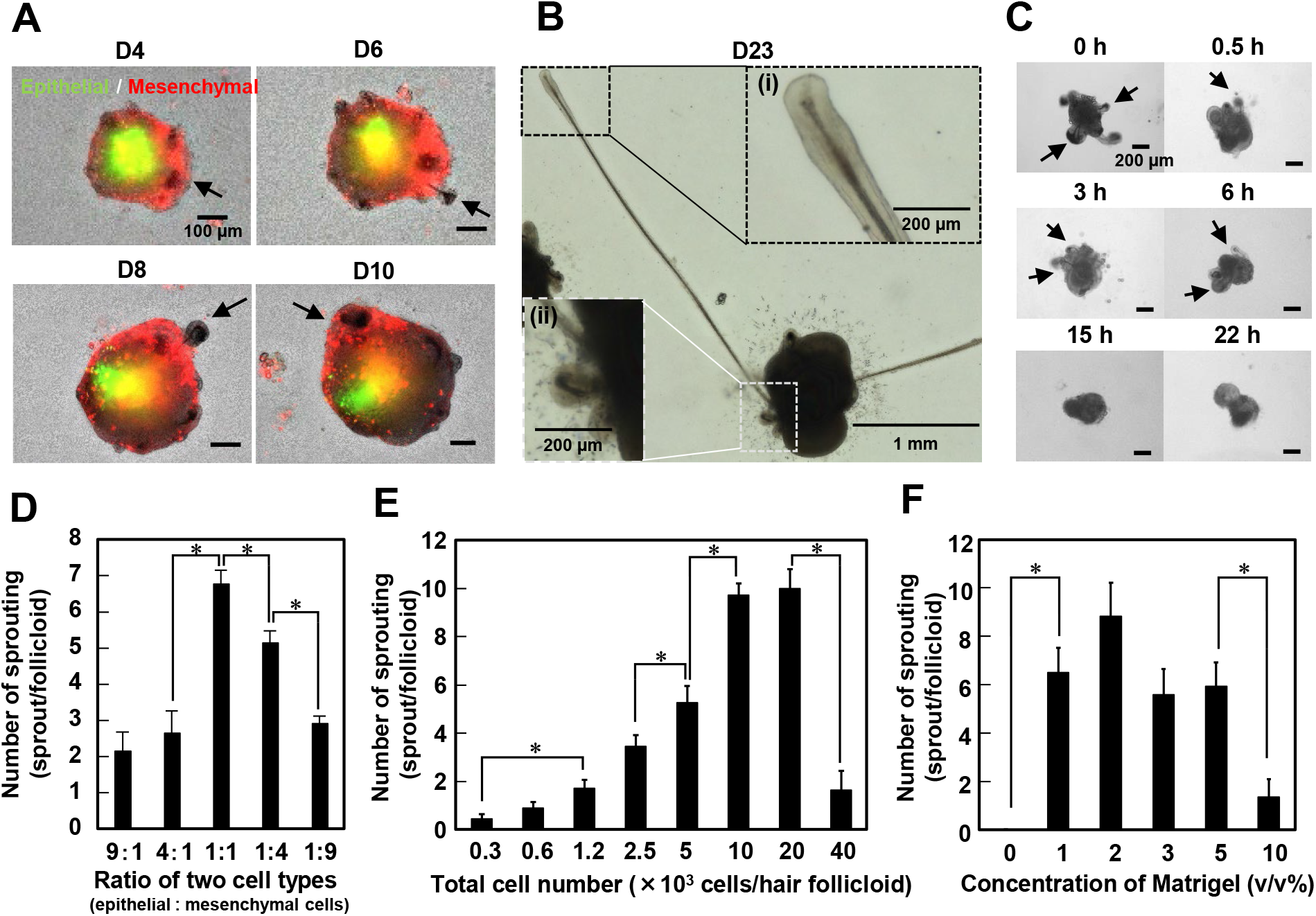
Characterization of hair follicloids. (**A**) Spatial distribution of epithelial and mesenchymal cells in hair follicloids. Epithelial and mesenchymal cells were fluorescently labeled with Vybrant DiO (green) and Vybrant Dil (red), respectively, prior to seeding. The two types of cells had separated and formed core-shell aggregates. Early hair follicle sprouting was observed at 4 days of culture (arrows). (**B**) A representative long sprouting hair follicle generated from hair follicloids after an extended period of culture. (i) Magnified views of the end of a papilla-like structure and (ii) a shorter and newly sprouted hair follicle. (**C**) Dependence of hair follicle sprouting on Matrigel supplementation at different time periods. A low concentration (2v/v%) of Matrigel was added at the indicated time points, and stereomicroscopic images were obtained at 8 days of culture. The arrows indicate hair follicle sprouting. (**D**) Dependence of hair follicle sprouting on the ratios of the two cell types. Hair follicles were counted at 8 days of culture. Numerical variables were statistically evaluated using Tukey’s tests; \**p* < 0.05, n = 20. (**E**) Dependence of hair follicle sprouting on the number of cells in a hair follicloid. Hair follicles were counted after 12 days of culture. Numerical variables were statistically evaluated using Tukey’s tests; *p < 0.05, n = 12. (**F**) Dependence of hair follicle sprouting on Matrigel concentrations. Hair follicles were counted after 12 days of culture. Numerical variables were statistically evaluated using Tukey’s tests; *p < 0.05, n = 12.

To further optimize the hair follicloid formation, several factors that may affect the hair follicle sprouting were investigated, such as the timing to add 2v/v% Matrigel, the mixing ratios of the two cell types, the total cell numbers, and the concentrations of Matrigel at the first step. Hair follicle sprouting was sensitive to the timing of Matrigel supplement (Fig. 2C). Vigorous sprouting was observed when the timing was 0–6 h after seeding, but when it was over 15 h, almost no sprouting was observed where the cells formed dumbbell-shape HFGs. The results were contrary to our expectations. Because cell aggregation is critical to induce EMIs, we expected that it would be better to add Matrigel after the cells were sufficiently aggregated as previously reported (*18*). However, Matrigel had to be supplemented before dense and compacted aggregate formation. These results imply that the involvement of Matrigel components such as extracellular matrices between the cells may be crucial to induce core-shell aggregates and subsequent hair follicle sprouting. In the following experiments, Matrigel was added immediately after cell seeding (0 h), and the culture was kept at 4 °C for 30 min. The mixing ratios of the two cell types significantly altered the sprouting (Fig. 2D). The sprouting efficiency peaked when the mixing ratio of epithelial and mesenchymal cells was equal, probably because EMIs might have been maximized. The sprouting efficiency was increased by increasing the total number of cells at the constant mixing ratio (epithelial and mesenchymal cells, 1:1) (Fig. 2E). However, at the highest cell number, the efficiency significantly dropped, which could be attributed to the shortage of oxygen and nutrients. There was no significant difference in the efficiency between 1 and 5v/v% of Matrigel, but Matrigel at 10v/v% formed a hydrogel in which most cells remained encapsulated as a single cell and did not form an aggregate, resulting in quite low sprouting efficiency (Fig. 2F). Based on these results, hair follicloids were prepared using 20 × 10^3^ total cells at a 1:1 mixing ratio by supplementing 2v/v% Matrigel with a cell suspension solution and subsequent 30-min cooling.

Next, we investigated the histology of the hair follicles in hair follicloids at 10 days of culture (Fig. 3A). Typical cell markers for the hair follicle and surrounding tissues were observed in the generated hair follicloids, including the adipose tissue marker oil red O; arrector pili muscles marker α-SMA (*19*); hair follicle stem cell markers CD34 (*20*) and Sox9 (*21*); basal keratinocyte marker K5 (*22*); proliferating cell (hair matrix cell) marker Ki67 (*23*); dermal papilla cell marker versican (*24*); melanosome markers gp100 (*25*) and TRP1 (*25*); and melanocyte stem cell marker c-Kit (*26*). The hair shafts generated possessed hair shaft-specific features, including the hair cuticle (Fig. 3B) and medulla (Fig. 3C). Hair cuticles and melanin granules were observed in the cross-sections of hair shafts similarly to those in the native hair shafts (Fig. 3D).

**Fig. 3.**
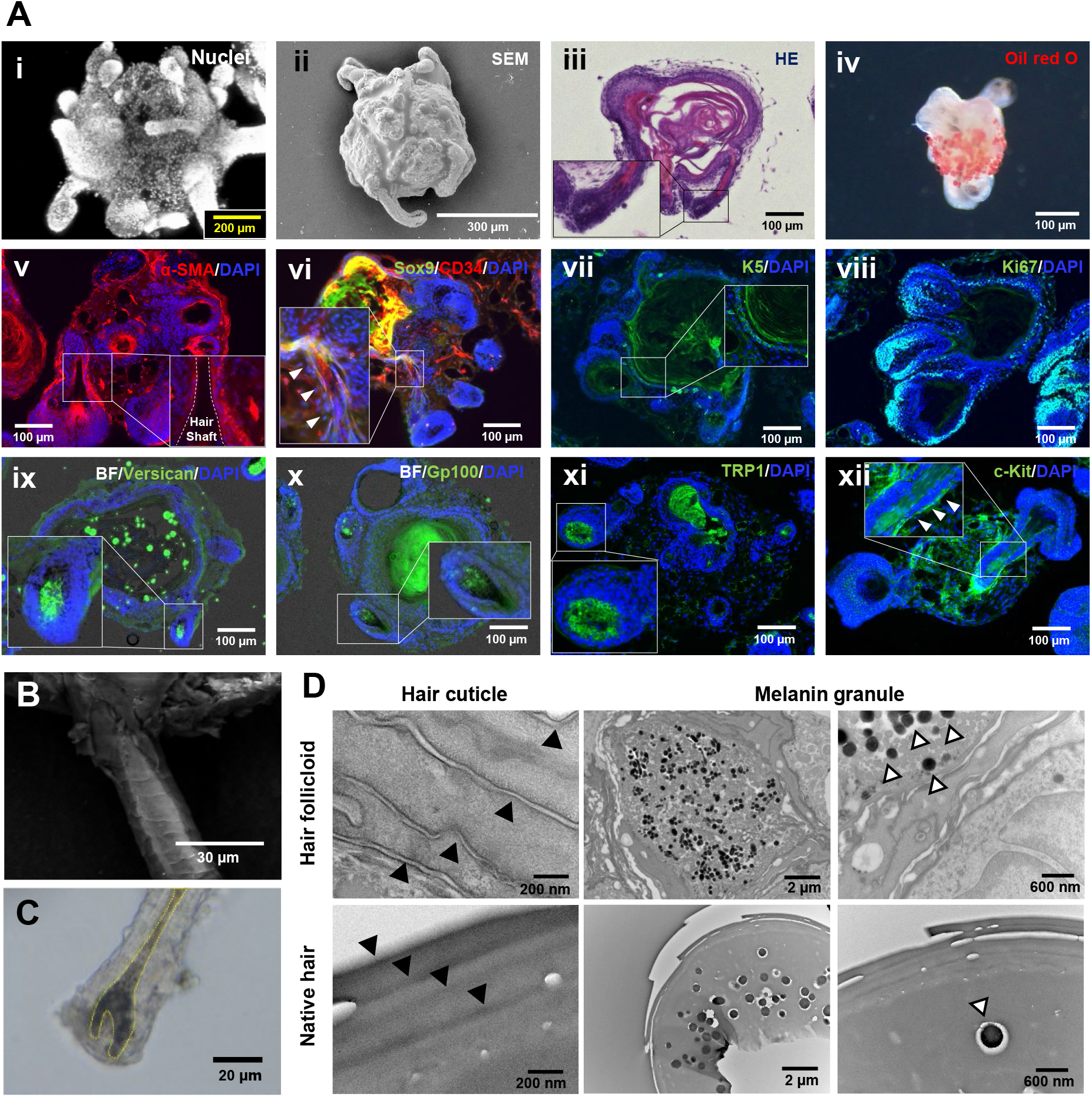
Analysis of hair follicloids and generated hair shafts. (**A**) Hair follicloids after 10 days of culture. Whole-mounted hair follicloids were stained with DAPI and observed using a confocal laser microscope (i) and scanning electron microscope (ii). The whole-mounted hair follicloids were stained with oil red O and observed using a stereomicroscope (iv). The hair follicloids were sectioned and stained with HE (iii) a fluorescent antibody against α-SMA (v), Sox9 (vi), CD34 (vi), K5 (vii), Ki67 (viii), versican (ix), gp100 (x), TRP1 (xi) and c-Kit (xii). The inserted images show magnified views of the box area in the images. BF: bright field. (**B**) Hair cuticles of generated hair shafts. The hair shaft was observed after 10 days of culture using a scanning electron microscopic. (**C**) Hair bulb of generated hair follicle after 10 days of culture. Hair follicloids were observed using a stereomicroscope. (**D**) Microstructures of generated and native hair shafts. Cross-sections were observed via transmission electron microscopy. White arrowheads indicate melanin granules and black arrowheads indicate hair cuticles.

### Genes involved in spatial arrangement of the two cell types in hair follicloids

The supplement of Matrigel at a low concentration significantly altered the spatial distribution of the two types of cells in the aggregate at 2 days of culture, as shown in Fig. 4A, and improved subsequent sprouting of hair follicles *in vitro*. We investigated the responsible molecular mechanisms to elucidate the link between the spatial cell arrangement and cell fate. We used the RT^2^ Profiler PCR array for ECM & adhesion molecules, which revealed that almost half the genes (39 out of 80 genes) were significantly upregulated, while one was downregulated in the presence of Matrigel compared with those without Matrigel (Fig. 4B). The genes upregulated >1.5-fold include fibulin-1 (*Fbln1*), fibronectin (*Fn1*), laminin (*Lama3, Lamb3, Lamc1*), collagen (*Col1a1, Col4a1, Col4a2, Col5a1*), versican (*Vcan*), transforming growth factor-beta-induced protein ig-h3 (*Tgfbi*), integrin β1 (*Itgb1*), and MMPs (*Mmp2, Mmp9, Mmp11, Mmp15*). Fibulin-1 is a secreted glycoprotein that binds to a number of ECMs including fibronectin, versican, and basal membrane proteins (*27*). TGFBI is a secreted RGD-containing protein that also binds to ECMs such as collagen types I, II, and IV as well as proteoglycans (*28*). MMP2 and 9 are gelatinase, of which main substrates are gelatin and collagen type IV (*29*). A furin cleavage site in the pro-peptide is a feature of membrane-bound MMPs including MMP15, but MMP11 is the secreted-type MMP that shares this feature, and both MMP11 and 15 degrade fibronectin and laminin (*30*). Collectively, the PCR array results imply that the hair follicloids reprogrammed their niches via the activation of the secretion and reconstitution of ECMs.

**Fig. 4.**
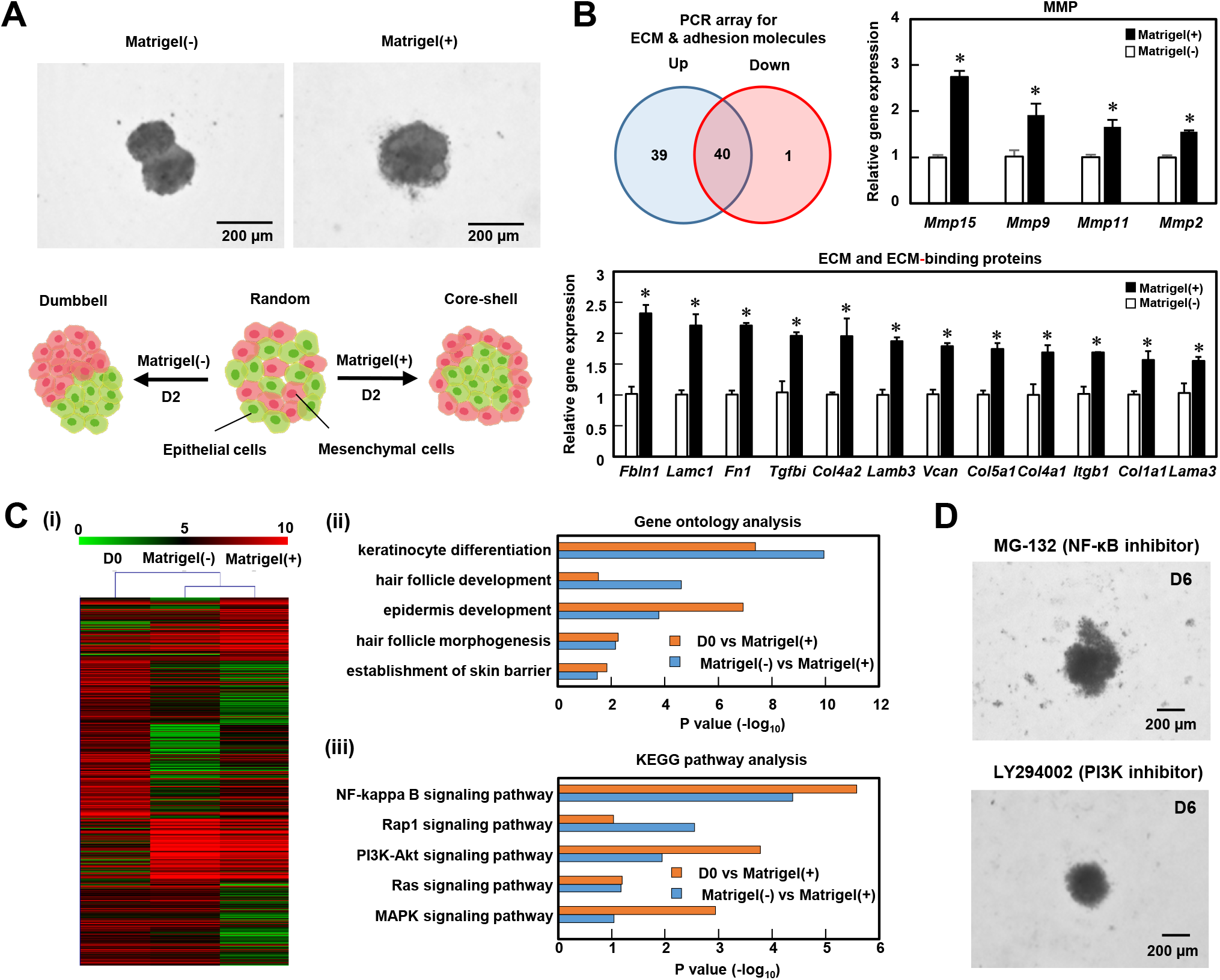
Gene expression analysis in hair follicloids. (**A**) Localization of epithelial and mesenchymal cells cultured with/without Matrigel supplementation after 2 days of culture. (**B**) The number of genes up-and down-regulated at 2 days of culture due to Matrigel supplementation. In total, 80 genes associated with extracellular matrices and cell adhesion molecules were examined. Numerical variables were statistically evaluated using standard *t*-tests and *p* < 0.05 was considered to be significant. The graph indicates the comparisons of three representative genes; *p < 0.05., n = 3. (**C**) DNA microarray analysis at 2 days of culture. Heat map of differentially expressed genes between Matrigel (+) and D0 as well as Matrigel (+) and Matrigel (-) (i). D0 indicates freshly isolated cells. Gene ontology analysis was performed for the genes associated with epidermal and hair development (ii). KEGG pathway analysis was performed for the upregulated, differentially expressed genes associated with hair follicle development (iii). (**D**) Effects of NF-kβ and PI3k pathway inhibitors on hair follicloid formation. Hair follicloids were cultured in a culture medium supplemented with MG-132, a NF-kappaβ inhibitor, or LY294002, a Pi3k inhibitor. Stereomicroscopic images of a hair follicloid were obtained after 6 days.

We performed global gene expression analysis to investigate involved gene sets and signal pathways. Transcription profiles were determined with a DNA microarray for the samples from a suspension of the epithelial and mesenchymal cells (D0), and at 2 days of culture with Matrigel (+) and without Matrigel (-) (Fig. 4C). Gene ontology analysis showed that genes associated with hair follicle and epidermis development were differentially expressed between D0 and Matrigel (+), or Matrigel (-) and Matrigel (+) (Fig. 4C-ii). Kyoto Encyclopedia of Genes and Genomes (KEGG) pathway analysis showed that the functioning of the NF-kappa B and PI3K-Akt signaling pathways was significantly different with both D0 vs Matrigel (+) and Matrigel (-) vs Matrigel (+) (Fig. 4C-iii). To investigate the role of the NF-kappa B and PI3K/Akt signaling pathways in *in vitro* hair follicle sprouting further, we performed targeted pathway inhibition using specific inhibitors. Supplementation with the NF-kappa B signaling antagonist, MG-132, and PI3K/Akt signaling antagonist, LY294002, completely suppressed hair follicle sprouting *in vitro* (Fig. 4D), indicating that both NF-kappa B and PI3K/Akt cascades are critical mechanism in hair follicle morphogenesis.

### Genes involved in hair follicle sprouting in hair follicloids

Wnt signaling is one of the fundamental mechanisms regulating embryonic development and is closely associated with hair follicle morphogenesis and differentiation (*31-33*). We investigated the expression of the Wnt signaling pathway in hair follicloids using the RT^2^ Profiler PCR array for Wnt Signaling Pathway. Note that the assay was performed at 4 days of culture when sprouting was initiated (Fig. 2A). The number of upregulated Wnt signaling-related genes were higher than those in the Matrigel (-) (Fig. 5A). Specifically, the expression levels of genes involved in hair follicle development, *Apc, Axin1, Lef, Lrp5*, and *Wnt10b* (*31, 34*), were significantly increased in hair follicloids (Fig. 5A and 5B).

**Fig. 5.**
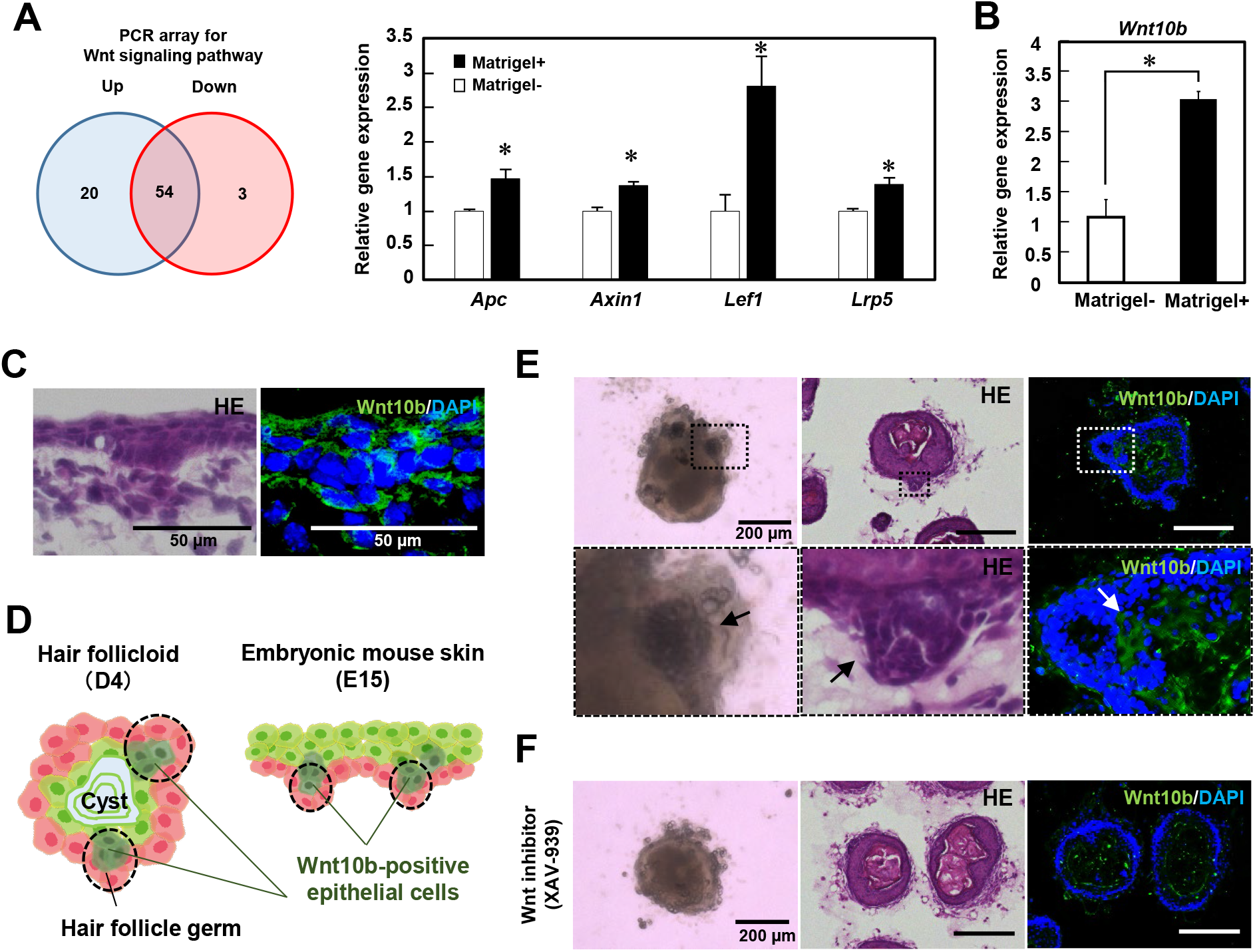
Wnt signaling in hair follicloids. (**A**) The number of genes up-and down-regulated because of Matrigel supplementation. A total of 77 genes associated with the Wnt signaling pathway were examined after 4 days of culture. Numerical variables were statistically evaluated using Student’s *t*-tests and p < 0.05 was considered to be significant. The graph shows a comparison of four representative genes; *p < 0.05, n = 3. (**B**) Wnt10b expression levels were assessed using RT-PCR. Numerical variables were statistically evaluated using Student’s *t*-tests; *p < 0.05, n = 3. (**C**) HE and immunohistochemical staining of sectioned embryonic mouse skin at E15. (**D**) Localization of Wnt10b positive cells in hair follicloids on day 4 of culture and E15 mouse skin. (**E**) Histological analysis of hair follicloids. Hair follicloids were sectioned and stained with HE and fluorescent antibodies against Wnt10b. The lower images show magnified views of the dash box in the upper images. The arrows indicate hair follicle germ-like structures. (**F**) Effects of the Wnt10b inhibitor on hair follicloid formation. Hair follicloids were cultured in the presence of XAV-939, a Wnt signaling inhibitor, for 4 days. Hair follicloids were sectioned and stained with HE and fluorescent antibodies against Wnt10b.

H&E and immunofluorescence staining revealed that HFG-like structures were generated in the outer layers of hair follicloids, and contained *Wnt10b* positive cells, as observed in the embryonic skin (Fig. 5C–E). XAV-939, a Wnt signaling antagonist, reportedly suppresses the expression of glycogen synthase kinase-3β (GSK-3β) and *Wnt10b* (*35*). The addition of XAV-939 to hair follicloids decreased the number of *Wnt10b* positive cells and inhibited the formation of HFG-like structures and subsequent hair follicle sprouting (Fig. 5F). We further visualized cells in the hair follicloid expressing the Wnt10b downstream molecule, Lef, using fluorescent reporter cells (Supplementary Fig. 2). These results suggest that the hair follicloids partially mimic normal embryonic hair folliculogenesis.

### Analysis of signal transduction for hair pigmentation in hair follicloids

In the hair follicloids, dynamics of melanosome transfer in hair bulbs were continuously monitored via time-lapse imaging using stereomicroscopy (Supplementary Movies 2, 3). Melanocytes are activated by α-melanocyte stimulating hormone (α-MSH) and produce melanosomes in the skin and hair follicles (*36*). We examined the effects of α-MSH on hair pigmentation (Fig. 6A). Hair follicloids treated with α-MSH generated dark-black hair shafts (Fig. 6B, Supplementary Movie 4) with an increased total melanin volume (Fig. 6C). Hair follicloids cultured in the presence of α-MSH exhibited an increased expression of four pigmentation genes, *TYRP1, TYRP2, TYR*, and *MITF*, at 6 days of culture (Fig. 6D). A mutation of melanosome transport-related genes including *Rab27a* causes hair graying through the so called Griscelli syndrome, a rare autosomal recessive disorder (*37*). Gray hair shafts were generated in hair follicloids when exposed to an antagonist of Rab27a, Nexinhib20 (Supplementary Fig. 3A–C). The volume of total melanin significantly decreased (Supplementary Fig. 3D) and the expression of pigmentation genes including TYRP1, TYRP2, TYR, and MITF decreased in the presence of Nexinhib20 (Supplementary Fig. 3e).

**Fig. 6.**
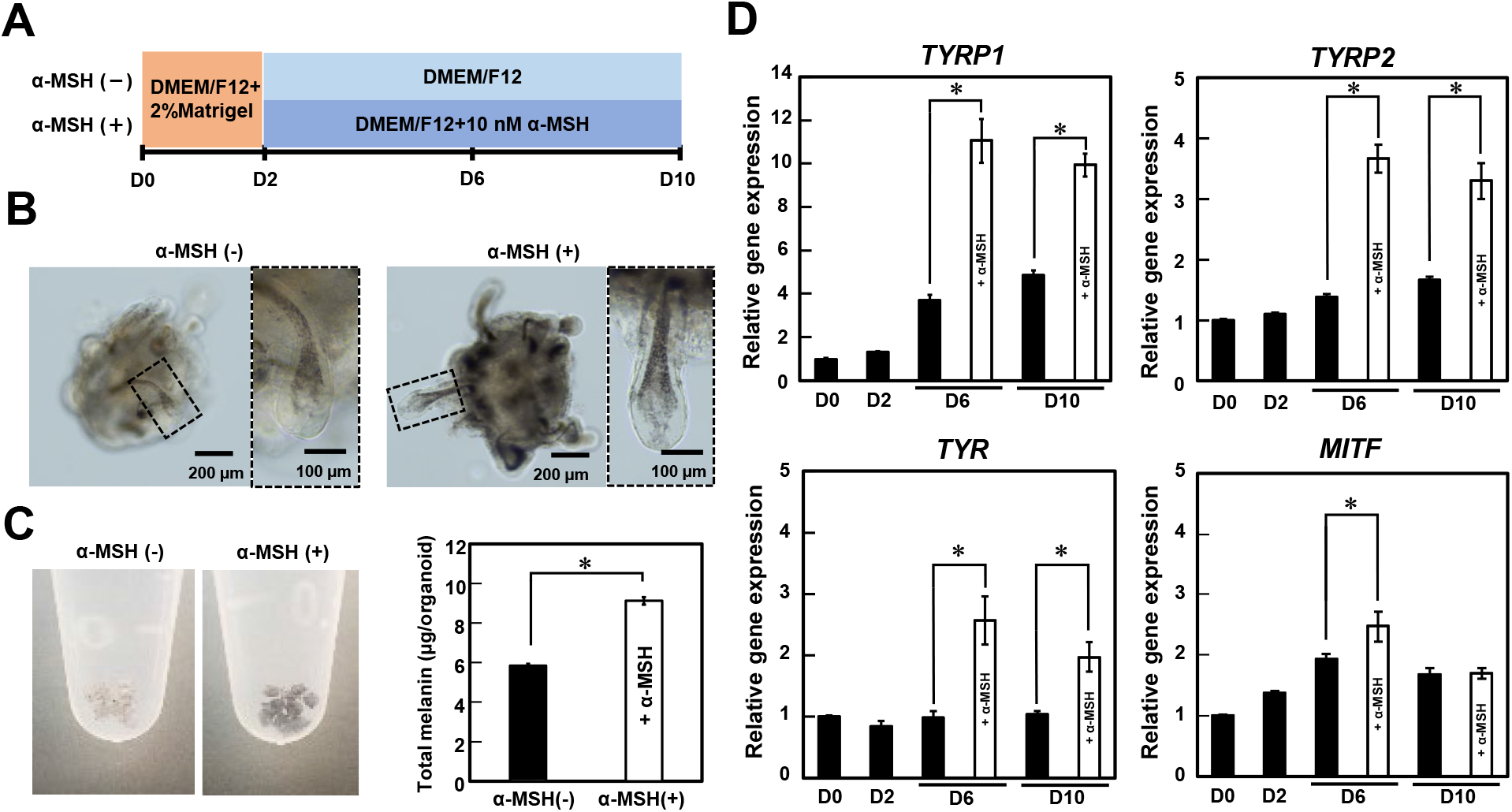
Effects of a hair pigmentation-related molecule on hair follicloids. (**A**) Procedures for testing the α-MSH pigmentation drug. (**B**) Microscopic images of hair follicloids cultured with/without α-MSH for 10 days. Hair follicloids were permeabilized and observed with a stereomicroscope. The images on the right show magnified views of the dash box in the images on the left. (**C**) Macroscopic views of hair follicloids cultured with/without α-MSH for 10 days. The graph shows the amount of melanin extracted from the hair follicloids. Error bars represent the standard error calculated from three independent experiments. Numerical variables were statistically evaluated using the Student’s *t*-test; *p < 0.05. (**D**) Gene expression levels associated with pigmentation. Error bars represent the standard error calculated from four independent experiments. Numerical variables were statistically evaluated using the Student’s *t*-test; *p < 0.05.

## Discussion

Organoid culture is a robust model for better understanding tissue development and maintenance since cell behavior during key events can be identified in a simplified and reproducible manner. Various tissue-specific organoids have been reported and utilized for the identification of disease-specific genes and novel signaling mechanisms responsible for regulating early embryonic development (*38, 39*). In the present study, we fabricated hair follicle organoids by controlling the spatial arrangement of epithelial and mesenchymal cells using quite a low concentration of Matrigel, resulting in highly efficient (∼100%) generation of hair shafts *in vitro*. Notably, the length of the generated hair shafts reached ∼3 mm after 23 days, which has not been achieved previously (*11, 40*). Because EMIs are crucial for the morphogenesis of other tissues and organs, this may provide a versatile approach for the preparation of other organoids.

Matrigel has been widely used in organoid research, and consists of basement membrane extracts from mouse sarcoma, including laminin, collagen IV, heparin sulfate proteoglycans, entactin/nidogen, and a number of growth factors (*41*). ECM extracts from the fetal skin promotes hair regeneration after transplantation with epithelial and mesenchymal cells (*42*). We aimed to identify components essential for the formation of hair follicloids. Hence, we examined laminin, collagen IV, entactin, growth factor reduced Matrigel, and collagen I and fibronectin on the hair follicloid culture. The results revealed that highly efficient hair shaft sprouting occurs using collagen IV, laminin-entactin complex, growth factor reduced Matrigel, and collagen I, suggesting that growth factors and laminin, the major component of Matrigel, are not directly involved in the hair follicle sprouting in the hair follicloids (Supplementary Fig. 4). It should be noted that collagen I, the most abundant ECM in the body, resulted in hair follicle sprouting at ∼100% efficiency, which would provide a robust, stable, and affordable system.

The transcriptome analysis of the hair follicloids revealed that signal pathways such as NF-kappaB and PI3K-Akt pathway were upregulated in the presence of either Matrigel (Fig. 4C) or collagen I (Supplementary Fig. 5), which is consistent with previous studies reporting that these genes are crucial for the early development of hair follicles and *de novo* regeneration of hair follicles (*43, 44*). Because hair follicle sprouting was virtually independent of the ECM types used in this study, the involvement of physicochemical environments, such as the viscosity of culture medium, in the formation of hair follicloids was also investigated. Culture media with different viscosities were prepared by adding methylcellulose, a cell inert polymer, and tested for hair follicloid formation, but no hair follicle sprouting was observed (data not shown). The use of cell-adhesive synthetic polymers, such as polyethylene glycol and polycaprolactone conjugated with cell adhesion motifs (*45*), can provide a more well-regulated organoid system, which will be the subject of our future research.

The hair follicle sprouting with Matrigel and the other ECMs was closely associated with the formation of core-shell-shape configuration rather than dumbbell-shape (Fig. 7). The mechanism that determines the spatial distribution of the two cell types may be explained by the differential adhesion hypothesis governed by interfacial energies (*46, 47*). The hypothesis states that an aggregate configuration composed of two cell types depends on the balance of cell adhesivity between homogeneous and heterogeneous cell types and eventually reaches to minimize the energy in the system. Monte Carlo simulations based on the hypothesis revealed that two types of cells separated from each other and eventually formed a dumbbell-shape aggregate when the adhesivity between heterogeneous cell types is weak compared with that between homogeneous cell types. In contrast, when adhesivity between the heterogeneous cell types is comparable to that between homogeneous cell types, the cells form an aggregate composed of more adhesive cells at the core and fewer adhesive cells in the outer layer. Our results imply that the supplement of ECMs increases the adhesivity between heterogeneous cell types (probably through integrin) to form a core-shell aggregate, which increases the contact area between two cell regions to enhance EMIs.

**Fig. 7.**
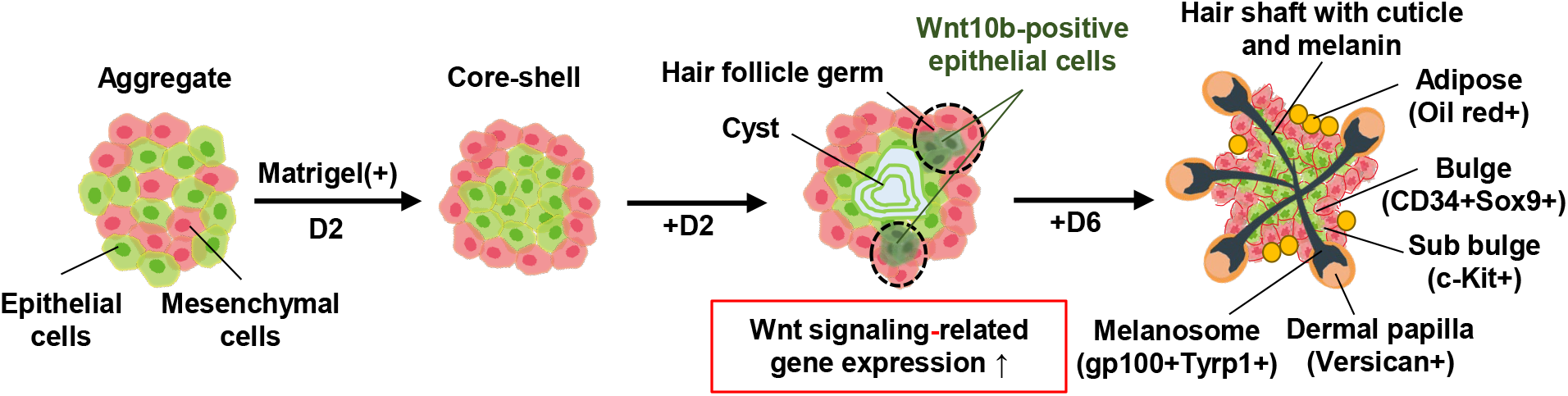
Schematic illustration of the process of hair follicloid formation. Embryonic epithelial and mesenchymal cells assembled via random distribution into a single aggregate. In the presence of Matrigel (non-gel-forming), the two types of cells separated from each other and self-organized into a core-shell aggregate within 2 days of culture; following which the aggregates generated epithelial cysts in the core and hair follicle germ-like structures containing *Wnt10b*-positive epithelial cells in the outer layer. After an additional 6 days of culture, hair follicle sprouting occurred and hair shafts growing from hair follicle germs became elongated. These hair follicles had specific features, including bulges, sub-bulge, dermal papillae, melanosomes, adipose tissues, and cuticles.

The global gene expression analysis showed that the remodeling of the ECMs seen in in *vivo* hair follicles (*48*), including laminins, fibronectins, and collagens, was activated in the hair follicloids (Fig. 4B). An approach has been reported to fabricate core-shell aggregates by seeding mesenchymal cells to form aggregates and by subsequently seeding epithelial cells to wrap the core aggregates (*49*). In accordance with this report, we prepared core-shell aggregates through two-step seeding, in which a core of epithelial cells was formed followed by the formation of a shell of mesenchymal cells. No hair follicle sprouting occurred with this approach, neither with nor without low concentrations of Matrigel (data not shown). These results indicate that the secretion of ECMs and their remodeling to form niches during the spontaneous tissue formation may be critical for hair follicle sprouting.

This approach is scalable, because hair follicloids are formed through the self-organization of cells when the two types of cells are aggregated in culture medium supplemented with a low concentration of ECM. In our previous study, we showed that a microdevice with an array of 1.0 mm diameter round-bottom wells made of oxygen-permeable silicone rubber could produce hundreds to thousands of dumbbell-shaped hair follicle germs under improved oxygen concentration conditions (*14*). Indeed, successful hair follicloid formation was observed in the microdevice in 9 days of culture (Supplementary Fig. 6).

The large-scale preparation of hair follicloids can be combined with genetic engineering technology (using CRISPR-Cas9, siRNA and signal inhibitor) to perform a comprehensive analysis of key genes related to hair follicle development, hair pigmentation, and hair follicle diseases. For example, crucial genes related to hair pigmentation (more than 650 genes) have been determined using transgenic mice with abnormal body hairs (*50*). However, laborious and time-consuming animal experiments and the lack of reliable *in vitro* models have hindered further elucidation. The hair follicloids may open a new avenue for such comprehensive studies.

In summary, we proposed an organoid culture system that generates hair follicles and hair shafts with almost 100% efficiency. This approach may be useful not only for understanding of the basis of EMIs in hair follicle induction but also for applications as alternatives to animal testing, hair follicle regeneration, and drug screenings. The limitation of hair follicloids is the lack of hair cycles. Hair loss disorder such as androgenic alopecia occurs due to the shortening of hair cycles. Recapitulation of hair cycles and such pathological nature *in vitro* should be preferable for understanding hair loss disorders and finding new drugs. In this study, rodent cells were used due to their shorter period of hair generation, higher hair inducing activity, and availability compared to human cells from hair follicle donors. The organoid culture system must be optimized with human cells.

Although it is challenging to culture human hair follicle epithelial stem cells and dermal papilla cells with current technology, recent substantial advances in molecular biology techniques, such as small molecule-induced chemical reprogramming or genetic reprogramming of genes, may provide potential cell sources.

## Materials and Methods

### Animals

Pregnant C57BL/6J mice were purchased from CLEA (Tokyo, Japan). The committee on animal care and use at Yokohama National University (Permit Number: 2019-04 and 2019-06), Central Institute for Experimental Animals (Permit Number: 20077) and National Institute of Health Sciences (Permit Number: 782) approved the animal study. The care and handling of mice conformed with the requirements of the committee on animal care and use at Yokohama National University, Central Institute for Experimental Animals and National Institute of Health Sciences.

### Preparation of murine epithelial cells and mesenchymal cells

Mouse epithelial cells and mesenchymal cells were made to dissociate from each other using a previously described method (*14, 51*). Briefly, the dorsal skin of mouse embryos (E18) was harvested under a dissection microscope and aseptically treated with 4.8 U/mL dispase II (Merck Millipore, Burlington, MA, USA) for 60 min at 4 °C, and epithelial and mesenchymal layers were separated using two pairs of tweezers. The epithelial layer was then treated with 100 U/mL collagenase type I (Fujifilm Wako Pure Chemical Corporation, Tokyo, Japan) for 80 min at 37 °C, followed by 0.25% trypsin for 10 min at 37 °C. The dermal layer was treated with 100 U/mL collagenase type I for 80 min at 37 °C. Debris and tissue aggregates were removed using a cell strainer (40 µm mesh, Corning, NY, USA). After centrifugation at 180 × *g* for 3 min, the epithelial and mesenchymal cells were suspended in DMEM/F-12 medium, which was prepared using Advanced Dulbecco’s Modified Eagle Medium/Ham’s F-12 (Thermo Fisher Scientific, Waltham, MA, USA) containing 1% GlutaMAX-I (Thermo Fisher Scientific) and 0.2% Normocin (InvivoGen, San Diego, CA, USA).

### Preparation of hair follicloids

To engineer hair follicloids, epithelial and mesenchymal cells were suspended in 0.2 mL advanced DMEM/F-12 containing 2 v/v% Matrigel (Corning) and seeded into the wells of a non-cell-adhesive, round-bottomed, 96-well plate (Prime Surface 96U plate, Sumitomo Bakelite, Akita, Japan). The plates were cooled at 4 °C in a freezer for at least 30 min and then incubated at 37 °C under 5% CO_2_ conditions. Cells were maintained in DMEM/F-12 and 0.1 mL of medium was exchanged for each well every 2 days. To generate longer hair shafts, hair follicloids cultured in 96-well plates for 3 days were embedded in Matrigel and cultured for an additional 20 days.

Hair shafts from hair follicloids were characterized using an all-in-one microscope (BZ-X810, Keyence, Osaka, Japan). The hair follicloid structures were analyzed via hematoxylin and eosin and immunohistochemical staining. The morphological features of the generated hair shafts, including cuticles, were examined using a scanning electron microscope (Miniscope, Hitachi High-Tech, Tokyo, Japan) at 10–15 kV. We examined cross-sections of hair shafts using a transmitted electron microscope.

### Inhibition of signaling pathways

To investigate the effects of NF-kappaβ, PI3K, and Wnt signaling inhibition on hair shaft generation, hair follicloids were cultured with DMEM/F12 medium containing 3 µM MG-132 (Abcam, Cambridge, UK), 50 µM LY294002 (Abcam), or 10 nM XAV-939 (Chem Scene, NJ, USA), respectively. Hair shaft generation was observed in the presence of MG-132 and LY294002 after 6 days. HFG formation and Wnt10b activation were evaluated via HE and immunohistochemical staining after 4 days.

### Microarray analysis

The total RNA content was extracted using a RNeasy mini kit (QIAGEN) from three samples (D0, epithelial and mesenchymal cell suspension; Matrigel (+), hair follicloids cultured for 2 days in the presence of Matrigel; Matrigel (-), hair follicloids cultured for 2 days in the absence of Matrigel). GeneChip analysis was performed by Takara Bio (Shiga, Japan). Briefly, isolated total RNA was reverse-transcribed and biotin-labeled using the GeneChip 3’ IVT PLUS Reagent Kit (Applied Biosystems, Waltham, MA, USA). Next, each labeled cRNA sample was fragmented and hybridized to the GeneChip Mouse Genome 430 2.0 Array (Applied Biosystems) and scanned using the GeneChip Scanner 3000 7G (Applied Biosystems) according to the manufacturer’s protocol. For microarray data analysis, normalized expression ratios were calculated for each gene and tested for significance, using the criteria that the expression ratio should be either > 2-fold or < 0.5-fold. The hierarchical clustering of differentially expressed genes was performed using MeV software. During analysis, genes exhibiting significant differential expression in both D2_Matrigel (+) versus D0 and in D2_Matrigel (+) versus D2_Matrigel (-) were selected. Gene ontology and KEGG pathway analysis were carried out by the gene batch viewer in DAVID (http://david.abcc.ncifcrf.gov/). During analysis, genes exhibiting significant differential expression in D2_Matrigel (+) versus D0 or in D2_Matrigel (+) versus D2_Matrigel (-) were selected.

The most enriched pathways associated with hair follicle and epidermis development were selected from “Gene Ontology” and “Pathways” and had significant P-values (p<0.05).

### Treatment with pigmentation agents

To investigate the effect of pigmentation drugs on the hair color of generated shafts, the DMEM/F12 medium was supplemented with 10 nM α-MSH (Merck Millipore, Burlington, MA, USA) on day 2 after seeding. Then, one-half of the DMEM/F12 medium containing 10 nM α-MSH was changed every 2 days. The hair shaft color was observed using an all-in-one microscope and digital camera, and the relative expression levels of genes associated with hair pigmentation were evaluated via real-time reverse transcription polymerase chain reaction (RT-PCR) analysis. The total amount of generated melanin was quantified using a plate reader (EnSpire™ Alpha, Perkin Elmer, Waltham, MA, USA).

### Histological and immunohistochemical staining

To perform histological staining, samples were washed with PBS, fixed with 4% formaldehyde overnight at 25 °C, rinsed 3 times with PBS, and successively submerged into 10%, 20%, and 30% sucrose solutions (FUJIFILM Wako Laboratory Chemicals, Osaka, Japan) for 1 h at 25 °C.

Next, the samples were transferred into a cryo-dish, the sucrose solution was carefully aspirated to ensure that spheroids alone would remain, and the cryo-dish was filled with the Tissue-Tek O.C.T. compound (Sakura Finetek, Tokyo, Japan). After embedding samples in the O.C.T. compound, 10-μm-thick sections were cut and stained using Meyer’s hematoxylin and eosin Y solutions.

For immunohistochemical staining, the 10-μm-thick frozen sections were prepared as described above. The samples were first blocked in PBS containing 10% BSA for 1 h at room temperature and subsequently incubated overnight with anti-Wnt10b (ab70816, Abcam), anti-melanoma gp100 (ab137078, Abcam), anti-α-SMA (ab7817, Abcam), anti-Sox9 (AB5535, Merck Millipore), anti-K5 (ab53121, Abcam), anti-Ki67 (ab16667, Abcam), anti-TRP1 (ab178676, Abcam), anti-c-Kit (ab273119, Abcam), and anti-versican (AB1033, Merck Millipore) at 4 °C. The samples were incubated with the corresponding secondary antibodies (ab150077, Abcam; A10037, A11078, A48270, Invitrogen) in the blocking solution for 2 h at 25 °C, in the blocking solution for 2 h at room temperature, and finally with DAPI in PBS for 10 min. A confocal microscope (LSM 700, Carl Zeiss, Germany) and all-in-one microscope (Keyence) were used for fluorescence imaging.

For oil red O staining, fresh whole aggregates were stained using oil red O staining kit (ScyTec). Briefly, samples were treated with propylene glycol for 5 min at 25 °C, oil red O solution for 10 min at 60 °C, propylene glycol solution diluted at a ratio of 17:3 in 1× PBS for 1 min at 25 °C, and distilled water for 1 min at 25 °C. Samples were imaged using an inverted microscope (CKX53, Olympus).

### Gene expression analysis using RT-PCR

Total RNA was extracted from samples using a RNeasy mini kit (QIAGEN), and cDNA was synthesized via reverse transcription using the RT^2^ First standard kit (QIAGEN) or ReverTra Ace™ RT-qPCR kit (Toyobo, Osaka, Japan), according to the manufacturer’s instructions.

Gene expression levels related to ECM and adhesion molecules and Wnt signaling proteins were identified using the RT^2^ Profiler PCR Array with Mouse Extracellular Matrix & Adhesion Molecules (PAMM-013ZA, QIAGEN) and the mouse WNT signaling pathway RT^2^ Profiler PCR array (PAMM-043Z, QIAGEN). We performed qPCR using the StepOne Plus RT-PCR system (Applied Biosystems) and TB Green^®^ Premix Ex Taq II reagent (Takara Bio). We used primers (Supplementary Table 1) to assess the expression levels of the following genes: *WNT10B, TYRP1* (Tyrosinase-related protein 1), *TYRP2* (Tyrosinase-related protein 2), *TYR* (tyrosinase), *MITF* (Microphthalmia-associated transcription factor), *RAB27A*, and *ACTB* (β-actin). The expression levels of all the genes were normalized to those of *GAPDH* (for gene expression related to ECM and adhesion molecules) or *ACTB*. Relative gene expression levels were determined using the 2^−ΔΔCt^ method and presented as the mean ± standard error values of at least three independent experiments.

### Measurement of the amount of melanin

To measure the level of intracellular melanin in hair follicloids, we added 100 μL of 1 N NaOH (Fujifilm Wako Pure Chemical Corporation) containing 10% DMSO to 10 hair organoids and heated the mixture at 80 °C for 90 min. The absorbance was then measured at 490 nm using a multimode plate reader (PE EnSpire alpha, Perkin Elmer). To derive the melanin levels from the absorbance values, we constructed a standard curve using absorbance values corresponding to a concentration range of 0–2.5 mg/mL of synthetic melanin (Sigma-Aldrich, St. Louis, MO) solution dissolved in 1 N NH_4_OH. We determined the average absorbance using at least three samples.

### Statistical analysis

Statistical analyses of the PCR array, RT-PCR, and total melanin volume data were conducted using two-tailed Student’s *t*-test, and results with *p* < 0.05 were considered significant. Since we anticipated variabilities between different donors, we repeated the experiments using at least three different donors. Statistical analyses for the optimization of the cell ratio, cell number, and Matrigel concentration were conducted using Tukey’s tests, and results with *p* < 0.05 were considered statistically significant. We cultured at least 12 hair follicloids per condition, which were used for the calculations of the number of hair follicles sprouting. Data are presented as the mean ± standard error.

## Supporting information

Supplementary materilals

## Author contributions

Conceptualization: TK, JF Methodology: TK, JF

Investigation: TK, AS, RA, RN, KS, YO

Visualization: TK, AS, RA, RN

Funding acquisition: TK, JF, YO

Writing: TK, JF

Resources: KS Supervision: JF

