## Supplementary materilals for "Reprogramming of three-dimensional microenvironments for *in vitro* hair follicle induction"

Supplementary Text

Long-term culture of hair follicle germs

To prepare hair follicle germs (HFGs), we suspended epithelial and mesenchymal cells (10 × 10^3^ cells/0.2 mL/well at 1:1 mesenchymal:epithelial cell ratio) in the DMEM/KG2 medium, which is a mixture containing DMEM and KG2 media (1:1 ratio), 10% FBS, and 1% penicillin-streptomycin. Suspensions containing the two cell types were mixed and seeded into the wells of a non-cell-adhesive, round-bottomed, 96-well or lab-made microarray plates. After 6, 15, and 23 days of culture, morphological changes in cell aggregates were examined using a phase contrast microscope (DP-71, Olympus, Tokyo, Japan), and the length of a generated hair shaft obtained from a cell aggregate surface was quantified from phase contrast images using ImageJ software (National Institutes of Health, Bethesda, MD, USA). The morphological features of generated hair shafts, including those of the hair cortex and microfibrils, were observed using a transmission electron microscope.

To evaluate the effects of FGF-2 on hair shaft generation, epithelial and mesenchymal cells were suspended in a culture medium containing 10 ng/mL of FGF-2 and seeded into a 96-well plate. After 10 days of culture, the number of hair strands generated *in vitro* was counted using a phase contrast microscope, and the relative expression of genes associated with hair follicle morphogenesis was evaluated via real-time reverse transcription polymerase chain reaction (RT-PCR) analysis.

Optimization of hair follicloid culture

To investigate the effects of Matrigel addition time on hair shaft generation, we supplemented the culture solution with Matrigel at different time points (after 0, 0.5, 3, 6, 15, and 22 h of culture) to ensure a 1:1 epithelial:mesenchymal cell ratio, 1×10^4^ of total cells, and 1% (v/v) of Matrigel. To investigate the effects of the ratio of two cell types on hair shaft generation, we prepared five solutions containing hair follicloids, with a 9:1, 4:1, 1:1, 1:4, or 1:9 epithelial: mesenchymal cell ratio, comprising a total of 1×10^4^ cells and 2% (v/v) of Matrigel. To investigate the effects of the total cell number of the two cell types on hair shaft generation, we prepared different solutions containing hair follicloids under eight different conditions. Thus, we prepared solutions with densities of 0.3, 0.6, 1.2, 2.5, 5, 10, 20, 40 ×10^4^ cells/0.2 mL, which included a 1:1 epithelial:mesenchymal cell ratio and 2% (v/v) of Matrigel. To investigate the effects of Matrigel concentrations on hair shaft generation, we prepared six Matrigel solutions containing hair follicloids with concentrations of 0%, 1%, 2%, 3%, 5%, and 10% (v/v) of Matrigel, comprising 1×10^4^ cells and 1:1 epithelial:mesenchymal cell ratio. To identify components of Matrigel that were effective for achieving hair shaft generation, we supplemented six components (1% laminin, 10% laminin, 1% laminin/entactin complex, 1 % collagen IV (all in v/v, from Corning [Corning, NY, USA]), 1% laminin/entactin complex + 1% collagen IV, and 1% growth factor reduced Matrigel) in DMEM/F-12 medium as a replacement for Matrigel. To prepare hair follicloids using type I collagen, we supplemented 2.4 mg/mL porcine type I collagen gel (Nitta gelatin, Morrisville, NC, USA) into 5% (v/v) in DMEM/F-12 medium as a replacement for Matrigel.

Large-scale preparation of hair follicloids

We fabricated microdevices via soft lithography processes. Briefly, microwell array configurations (diameter, 1 mm; pitch, 1.3 mm; depth, 1 mm; well number, 19 well) were designed and a mold was produced accordingly. A PDMS solution (consisting of a 10:1 mix of a pre-polymer solution and curing agent, Shin-Etsu Silicone, Tokyo, Japan) was poured onto the mold and cured in an oven at 80 °C. The thickness of the PDMS substrate at the floor of the microwells was set to 1 mm by adjusting the volume of utilized PDMS solution. The surface of the PDMS substrate, including the microwells, was modified via exposure (6–18 h) to a prevelex® solution (Nissan Chemical Corporation, Tokyo, Japan) to make it a non-cell-adhesive surface. To prepare multiple hair follicloids, cell suspensions (200 µL) containing epithelial and mesenchymal cells (total density 9.5 × 10^5^ cells/mL, 1:1 ratio) were poured onto the chips. The chips were cooled at 4 °C in a freezer for at least 30 min and then incubated at 37 °C in an incubator containing 5% CO_2_. Hair follicloids were observed after 9 days using an inverted microscope (CKX53, Olympus, Tokyo, Japan).

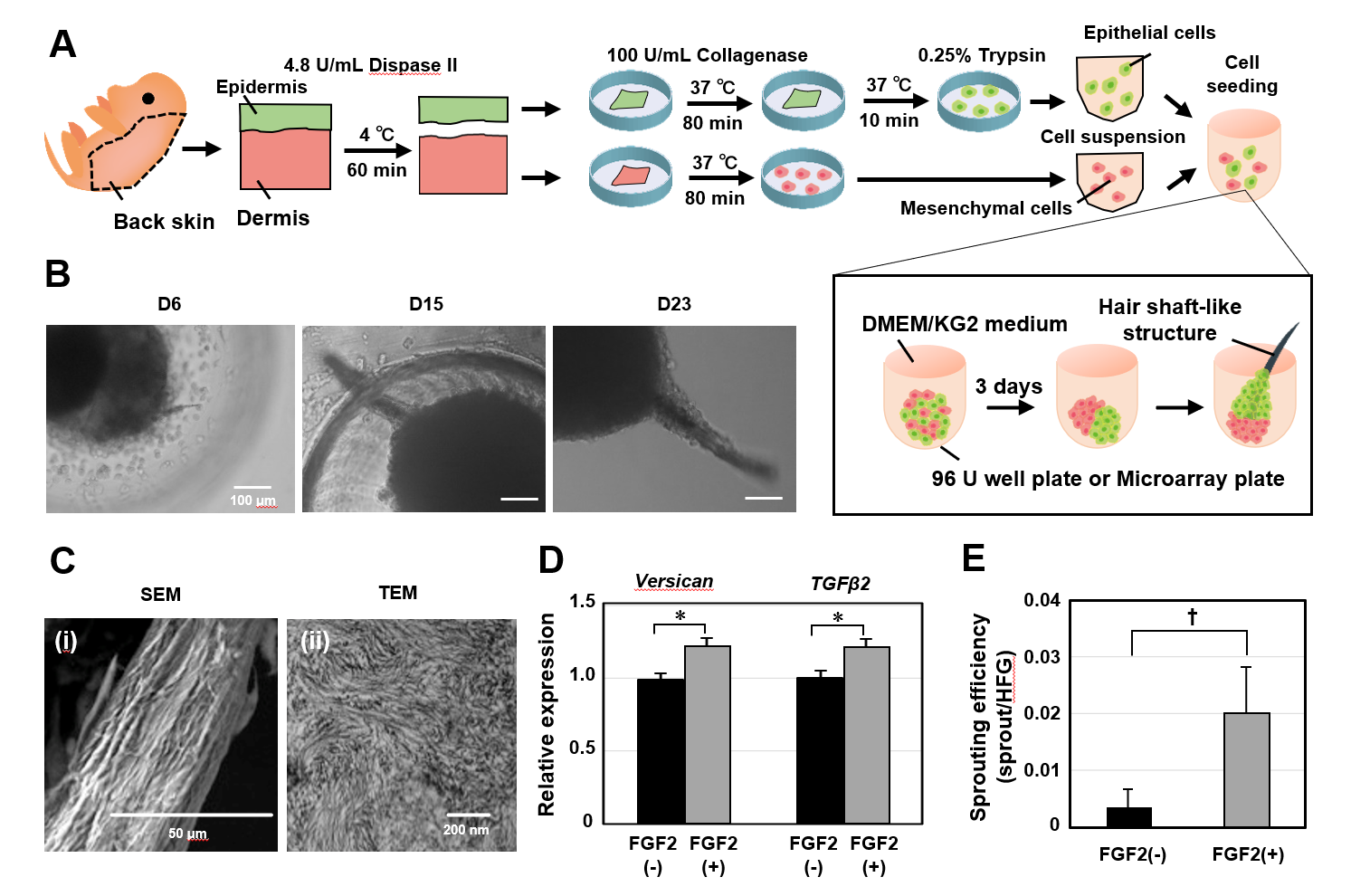

Fig. S1.

*In vitro* hair generation from self-sorted hair follicle germs. (a) Schematic representation of epithelial and mesenchymal cell isolation, spontaneous self-sorted hair follicle germ (ssHFG) formation, and hair shaft generation. (b) Phase-contrast microscopic images of a generated hair shaft sprouted from ssHFGs after 6 days of culture, which reached ~300 µm long after 23 days of culture. (c) Microstructures of a generated hair shaft. (i) Scanning electron microscopy and (ii) transmission electron microscopy revealed the morphological characteristics of the hair cortex and microfibrils. (d) Effects of FGF2 on the expression of hair induction genes, *Versican* and *TGFβ2*, evaluated after 10 days of culture. Error bars represent the standard error calculated from three independent experiments. Numerical variables were evaluated using the Student’s *t*-test; **p* < 0.05. (e) Effects of FGF2 on hair generation. The average amount of hair generated from FGF2-treated and non-treated ssHFGs was compared after 10 days of the seeding process. Error bars represent the standard error calculated from 300 independent experiments. Numerical variables were evaluated using the Student’s *t*-test; ^†^*p* < 0.1.

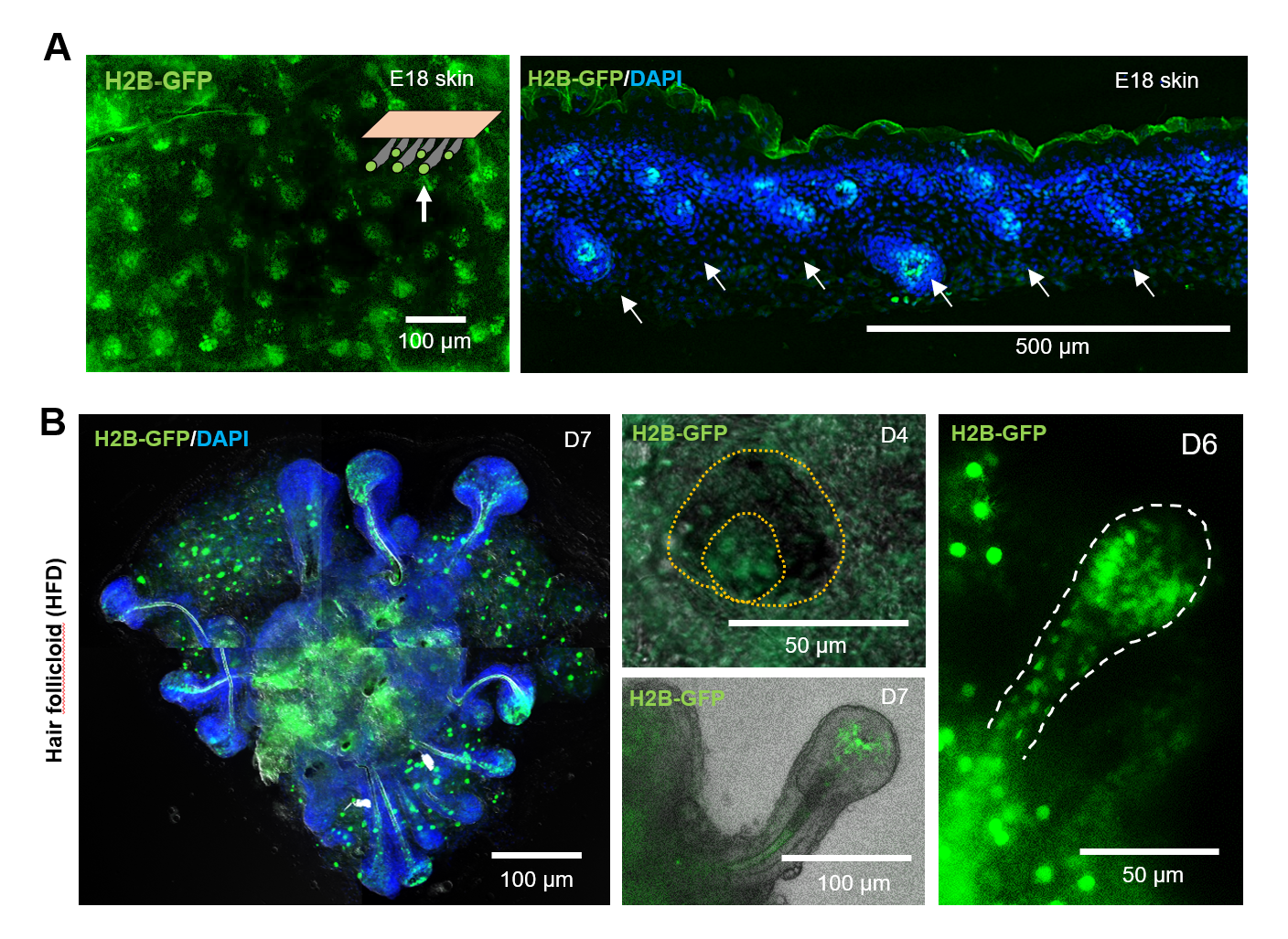

Fig. S2.

Wnt-βcatenin signaling in hair follicloids. (a) Localization of TCF/Lef-positive cells in E18 TCF/Lef:H2B/GFP mouse skin. (b) Localization of TCF/Lef positive cells in hair follicloids on days 4, 6, and 7 of culture.

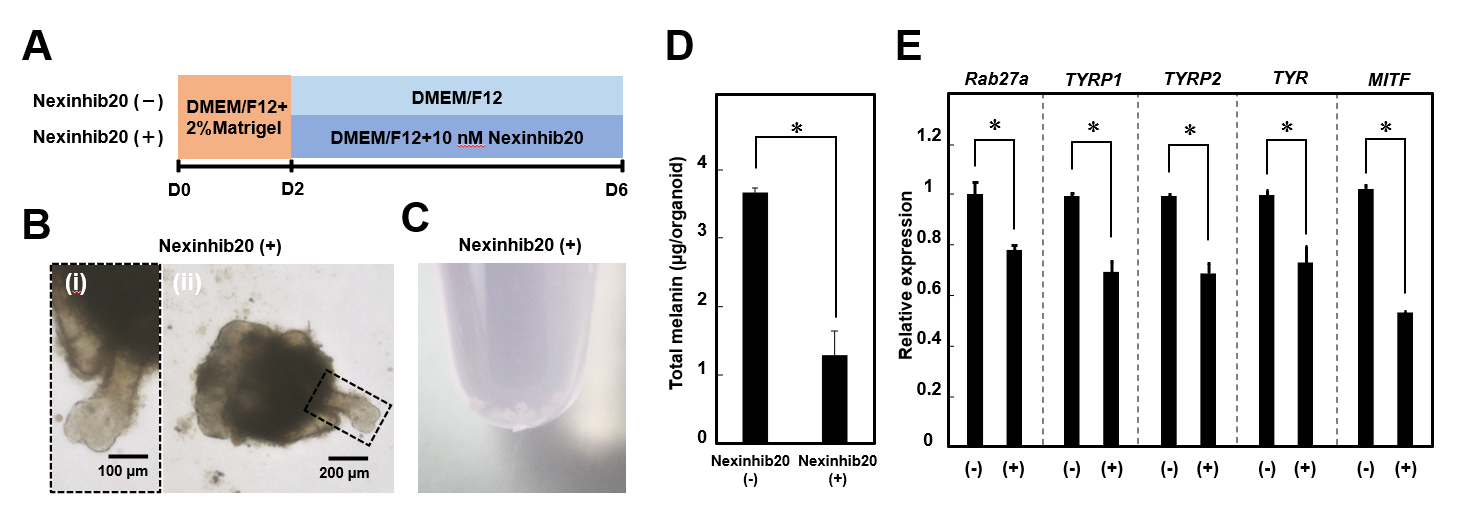

Fig. S3.

Effects of a depigmentation drug on hair follicloids. (a) Schematic representation of the methods used for screening of the depigmentation drug Nexinhib20. (b) Hair shafts generated from hair follicloids cultured with Nexinhib20 after 6 days of culture. Left panels (i) show a magnified view of the boxed areas in the right panels (ii). (c) Image of a hair follicloid cultured with Nexinhib20, obtained with a digital camera. (d) Melanin volumes in hair follicloids cultured with/without α-MSH after 6 days. Error bars represent the standard error calculated from three independent experiments. Numerical variables were evaluated using the Student’s *t*-test; **p* < 0.05. (e) Changes in gene expression-related hair pigmentation in cells from hair follicloids cultured with/without Nexinhib20. Error bars represent the standard error calculated from three independent experiments. Numerical variables were evaluated using the Student’s *t*-test; **p* < 0.05.

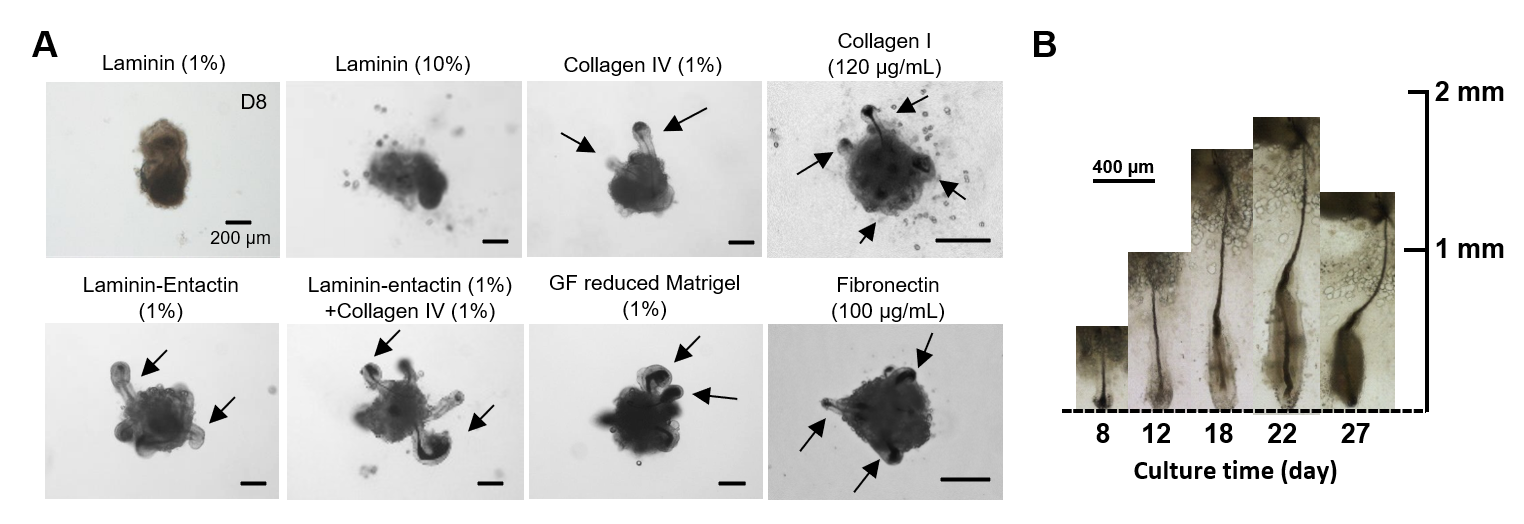

Fig. S4.

Effects of major/minor Matrigel components on hair follicle sprouting. (a) Hair follicle sprouting observed in components of Matrigel. Upon culturing cells in a laminin-supplemented medium, dumbbell-shaped cell structures were formed; no hair shaft generation occurred after 8 days of culture. In the presence of collagen IV and the laminin/entactin and laminin/entactin/collagen IV complexes, hair follicloids efficiently generated hair shafts. Replacement with growth factor-reduced Matrigel showed results comparable to those observed with Matrigel, suggesting that growth factors might not be crucial for hair follicle sprouting. The culture of hair follicloids with only minor constituents of Matrigel including collagen I and fibronectin efficiently resulted in hair follicle sprouting. Stereomicroscopic images were obtained after 8 days of culture. The arrows indicate hair follicle sprouting. (b) A representative long sprouting hair follicle generated from hair follicloids after an extended period of culture.

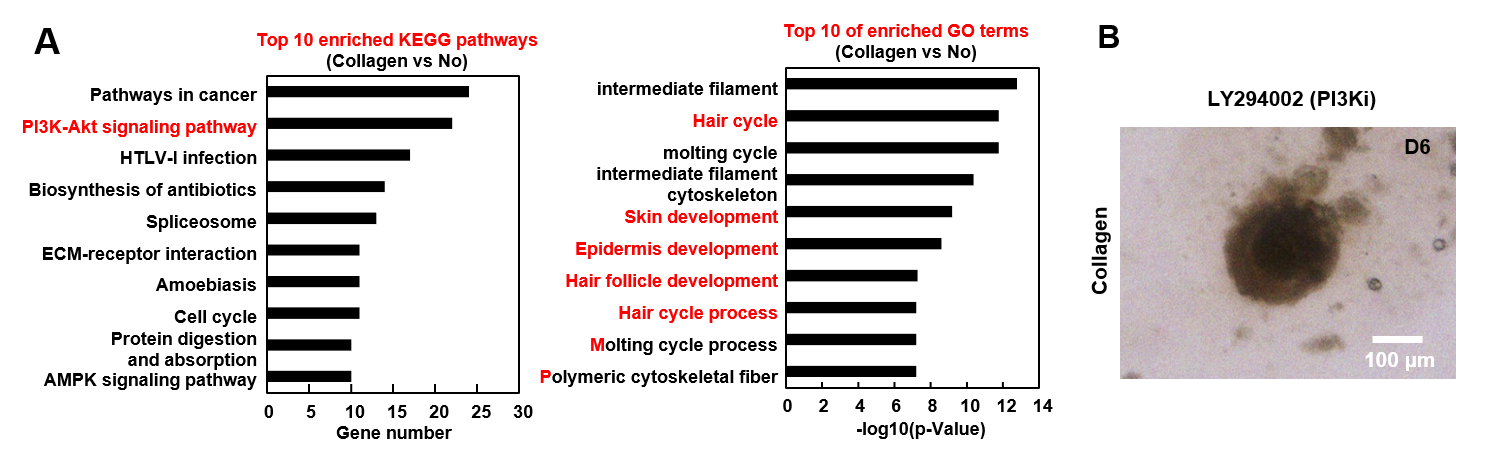

Fig. S5.

Gene expression analysis in hair follicloids in the presence of collagen. (a) DNA microarray analysis. The top 10 results for gene ontology (GO) and Kyoto Encyclopedia of Genes and Genomes (KEGG) pathway enrichment by genes with upregulated expression in hair follicloids in the presence of collagen as compared to that without collagen. (b) Effects of PI3k pathway inhibitor (LY294002) on hair follicloid formation. Hair follicloids were cultured in a medium supplemented with LY294002. Stereomicroscopic images of the hair follicloids were obtained after 6 days of culture.

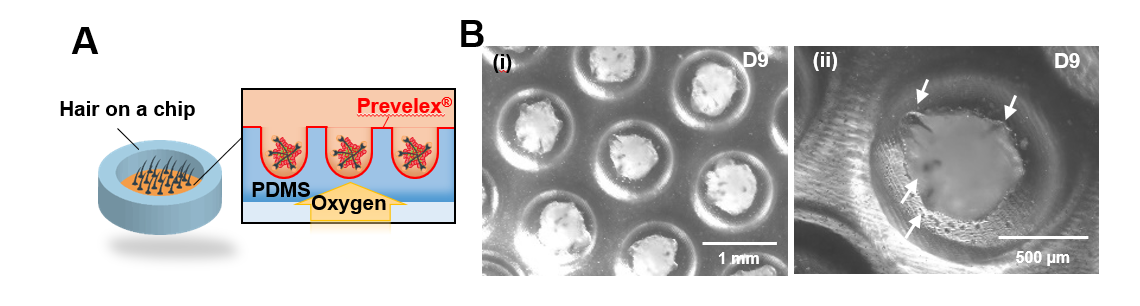

Fig. S6.

Large-scale preparation of hair follicloids. (a) Schematic representation of the large-scale preparation. Microfabricated microwell array plates composed of oxygen-permeable silicone elastomers were used for the large-scale preparation of hair follicloids. (b) Hair follicloids formed in a microfabricated microdevice for large-scale preparations. Stereomicroscopic images were obtained to visualize hair follicle sprouting (i, ii). The arrows indicate hair follicle sprouting.

Table S1.

List of primers used in the study

| Genes | Forward primer (5ʹ→3ʹ) | Reverse primer (5ʹ→3ʹ) |
| --- | --- | --- |
| *WNT10B* | CCAAGAGCCGGGCCCGAGTGA | AAGGGCGGAGGCCAGACCG |
| *TYRP1* | CGATACCCTGGGAACACT | TACACGGACCTCCAAGCA |
| *TYRP2* | CCAACGCTGATTAGTCGGA | GAAGAAGGGAGGGCTGTCA |
| *TYR* | AGCCTGTGCCTCCTCTAA | AGGAACCTCTGCCTGAAA |
| *MITF* | AGGACCTTGAAAACCGACAG | GTGGATGGGATAAGGGAAAG |
| *‎RAB27A* | CAAACAGCTTCCAGCTAAGGAC | GAGAACTCTGTGCCTCACCTCA |
| *ACTB* | TTGCTGACAGGATGCAGAAG | ACATCTGCTGGAAGGTGGAC |

Movie S1.

Sprouting of pigmented hair shafts generated from hair follicloids.

Movie S2.

Melanosome transport in a hair follicloid.

Movie S3.

Melanosome transport in a hair follicloid (high magnification).

Movie S4.

Sprouting of pigmented hair shafts generated from hair follicloids cultured with/without α-MSH.
